# Regulation of the Epstein-Barr Epigenome by Histone Variant MacroH2A

**DOI:** 10.64898/2026.11.11.765976

**Authors:** Leonardo Josué Castro-Muñoz, Leena Yoon, Bhanu Chandra Karisetty, Aradhana Bharti, Italo Tempera, Samantha S. Soldan, Paul Lieberman

## Abstract

Histone variants play an important role in regulating chromatin dynamics of Epstein-Barr Virus (EBV). Here, we investigate the role of histone variant macroH2A in EBV chromatin structure, gene expression and copy number control during latent infection. ChIP-seq analysis in EBV^+^ SNU719 gastric carcinoma and Mutu I Burkitt lymphoma cells revealed that macroH2A was most enriched at repetitive DNA elements, including oriLyt, BHLF1, LF3 and terminal repeat (TR) regions of the EBV genome. MacroH2A depletion in EBV-associated gastric cancer cell line SNU719 led to a small increase in some viral lytic genes and increase in viral DNA copy number. RNAseq transcriptomics after macroH2A depletion identified major perturbations in cellular gene pathways, including β-catenin/TCF signaling and chromatin remodeling. Depletion of macroH2A during EBV lytic reactivation reduced EBV lytic gene activation, but increased EBV DNA copy number. Previous study demonstrated a colocalization and cooperative binding between histone variant H2A.Z and EBNA1. We now show that depletion of macroH2A causes an increase in both H2A.Z and EBNA1 at oriP and cellular binding sites for EBNA1. We also found that γH2AX localizes to oriP and that this depends on macroH2A. Finally, we show that CRISPR/Cas9 knockout of macroH2A increases the replication and episome maintenance of oriP-containing plasmids. Taken together, these findings indicate that macroH2A localizes to the repetitive elements of EBV, including oriLyt, and can function to limit H2A.Z and EBNA1 binding at oriP, which alters DNA damage signaling at oriP and limits episomal replication. These findings provide a novel role for macroH2A in maintaining EBV copy number and genome integrity.

## 1 Introduction

Epstein-Barr virus (EBV) establishes long-term latent infection in over 90% of the adult population, and is causally linked to several malignancies, including endemic forms of Burkitt’s lymphoma and nasopharyngeal carcinoma [1; 2; 3] and a subtype of gastric carcinomas (GC) [4; 5]. During EBV latent infection the viral genome is maintained as multicopy, chromatin-associated episomes [6; 7]. Viral gene expression and DNA replication during latency is regulated by various epigenetic factors, including the incorporation of histone variants [8; 9]. Histone variants play an important role in the chromatin control of transcriptional regulation, DNA replication and DNA repair [10; 11]. How histone variants contribute to EBV latency and oncogenesis remains an area of significant interest.

EBV episome maintenance is maintained largely through the binding of the EBV encoded nuclear antigen EBNA1 to the viral origin of plasmid replication (oriP) [6; 7]. The chromatin organization of oriP is complex, subject to cell-cycle changes, and important for episome persistence, genome integrity and overall viral gene expression programming. We previously reported that the histone variant H2A.Z colocalizes with EBNA1 at oriP and other binding sites in the viral and host genomes [12]. H2A.Z depletion led to loss of EBNA1 binding at oriP and reactivation of EBV lytic gene expression and DNA replication. H2A.Z family are one of several histone H2A variants that include H2AX and macroH2A [13]. How these other histone H2A variants impact EBV latency and EBNA1 function is not yet known.

MacroH2A consists of two genetic isoforms (H2AYF1 and H2AFY2) that have both been implicated in numerous chromosome structures and control of various cellular processes ranging from X-chromosome inactivation to cancer [14; 15; 16; 17]. MacroH2As are unique among histone variants due to their C-terminal macro-domain. The macro-domain of macroH2A1.1 has been implicated in binding poly-ADPribose [18; 19], while other macrodomains may interact with RNA or other metabolites. Macro H2As have been found to regulate promoter-enhancer contacts [20], mediate X-chromosome inactivation through binding of chromatin remodeller ATRX [21], regulating heterochromatin through interaction with the histone chaperone ANP32B [22] and orchestrates synchronous replication origin firing on the inactive X-chromosome [23]. Additional biological functions attributed to macroH2As include inhibiting the EMT/MET transitions to maintain mesenchymal identity in pluripotent stem cells [24] and enforcing a dormancy phenotype of disseminated metastatic cancer cells by modulating gene programs in senescence and quiescence [25]. Thus macroH2As have been implicated in regulating repressive chromatin states critical for various normal and pathogenic biological processes.

Here we investigate the role of macroH2A in the control of EBV latency. We found that macroH2A is enriched at oriLyt and other repetitive elements, including non-coding RNA BHLF1 and LF3, and terminal repeats (TR). These genetic elements are of high biological significance since they are essential for viral latent-lytic switch and genome integrity [6]. We found the depletion of macroH2A increases the episome copy number through an increase in H2A.Z and EBNA1 binding at oriP. Our findings suggest that macroH2A antagonizes H2A.Z and EBNA1 binding at both viral and cellular sites to control EBV episome copy number in latently infected cells.

## 2 Methods

### Cell lines

SNU719, Mutu I, YCCEL1, C666-1 and AGS cells were grown in RPMI1640 media supplemented with 10% fetal bovine serum, 1X Glutamax, and 100 μg/ml streptomycin, and 100U/ml penicillin, cells were incubated at 37°C with 5% CO_2_ in a humidified chamber. 293T (ATCC) cells were maintained in Dulbecco’s modified Eagle’s medium (DMEM) supplemented with 10% fetal bovine serum, 100 μg/ml streptomycin, and 100U/ml penicillin. Cells were cultured in an incubator set at 37 °C and 5% CO_2_.

### Drug treatments

SNU179 cells were treated with Decitabine (DCB) at 7.5 µM, Ganciclovir (GSV) at 20 µM or an equivalent amount of dimethylsulfoxide (DMSO) for 72 hr. Cells were harvested and the Western blots and DNA copy numbers were performed.

### macroH2A knockdown by siRNA

SNU179 cells were transfected with small interfering RNAs targeting siH2AY (L2-011964-01-0020) or siControl (D2-001810-10-20) (Dharmacon, Lafayette, CO, USA) used DharmaFECT transfection reagents (Dharmacon, Lafayette, CO, USA). The cells were collected after 6 days and then subjected to further analyses.

### RNA extraction and cDNA synthesis

Total RNA was isolated from cells using the RNeasy Mini kit (Qiagen, Hilden, Germany), according to the manufacturer’s protocol. RNA was resuspended in 30 μL of RNAse-free H2O, which was treated with the DNAse Free DNA removal kit (Thermo Fisher Scientific, Waltham, MA, USA) and quantified. Reverse transcription was carried out on equal amounts of DNase-treated RNA using SuperScriptIV reverse transcriptase kit (Invitrogen), following the manufacturer’s instructions. qPCR was performed with Power SYBR green 2X PCR master (ThermoFisher Scientific, Waltham, MA, USA) mix with specific primers. The expression of EBV genes was verified by RT-qPCR using specific primers (**Table 1**) and normalized to cellular GUSB gene expression.

**Table 1.**
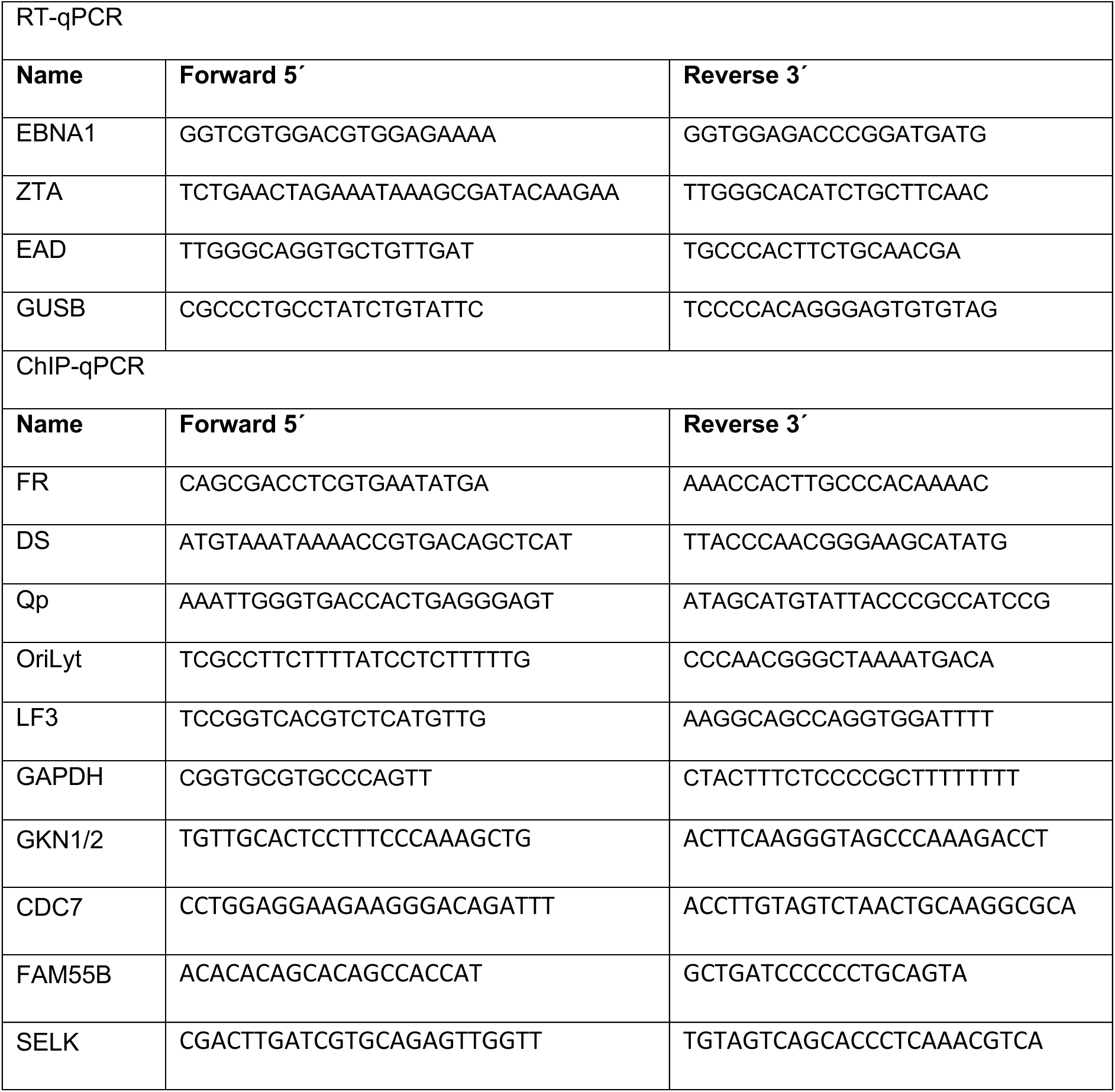
List of primers used.

### RNA-seq

Total RNA was isolated from each condition using the RNeasy Mini kit (Qiagen, Hilden, Germany), according to the manufacturer’s protocol. RNA was resuspended in 30 μL of RNAse-free H2O, which was treated with the DNAse DNA-Free DNA removal kit (Thermo Fisher Scientific, Waltham, MA, USA) and quantified by Qubit (Thermo Fisher Scientific, Waltham, MA, USA). RNA samples were either used for downstream RT-qPCR or submitted to the Wistar Institute genomics core facility for RNA quality control and sequencing library preparation using the SENSE mRNA-Seq Library Prep Kit V2 (Lexogen) to generate Illumina-compatible sequencing libraries according to the manufacturer’s instructions. Paired-end reads of 75 bp were obtained using a Illumina HiSeq 2500 sequencer. RNAseq data was aligned using STAR [26] against hg19 version of the Human genome and all unaligned reads were then aligned against NC_007605.1 version of EBV genome and RSEM v1.2.12 software was used to estimate raw read counts and RPKM for Human and EBV genes. Differential gene expression analysis was performed using DESeq2 R package [27], statistically significant differences in gene expression were determined using a threshold of p value< 0.05. Functional enrichment analyses were conducted using EnrichR [28] and Gene Set Enrichment Analysis (GSEA) [29] based on DESeq2 results with statistically significant genes. Gene Ontology (GO) enrichment analysis on biological and functional processes was then performed on the clusters. The dataset is deposited in Gene Expression Omnibus (GEO) under the accession number GSE.

### Western blot

Cells were collected and protein extracts obtained by adding Laemmli sample buffer. Equal amounts of proteins were resolved in 8–16% Novex Tris-Glycine gels (Invitrogen). Then transferred onto a Nitrocellulose membrane (BIO-RAD-1620112). The membrane was blocked using 8% milk in TBS/0.01% Tween buffer and then incubated with the indicated antibodies. Primary antibodies were prepared in TBS/0.01% Tween buffer as follows: rabbit monoclonal anti-macroH2A (diagenode-C15210003), rabbit polyclonal anti-EBNA1 (Pocono Rabbit Farm custom preparation), Anti-β-Actin (Sigma-A3854), and mouse monoclonal anti-ZTA (sc-53904-HRP) Membranes were washed three times with TBS/0.01% Tween buffer and incubated with HRP coupled secondary anti-mouse or anti-rabbit antibodies (Bio-Rad,). Finally, Proteins were visualized using the Immobilon Forte (Merck Millipore-WBLUF0500) according to the manufacturer’s instructions.

### Relative DNA copy number assay

Cells (1 × 10^6^ cells per sample) were collected and resuspended in 100 μl of SDS lysis buffer (1% SDS, 10 mM EDTA, 50 mM Tris [pH 8.0]). After 20 cycles of sonication (Diagenode Bioruptor), immunoprecipitation dilution buffer (0.01% SDS, 1.1% Triton X-100, 1.2 mM EDTA, 16.7 mM Tris [pH 8.0], 167 mM NaCl) was added to 1 ml, followed by incubation with proteinase K for 2 to 3 h at 50°C. Three hundred microliters of lysate were then removed and subjected to phenol-chloroform extraction and ethanol precipitation. Precipitated DNA was assayed by RT-qPCR. Relative EBV copy numbers were determined using primers (Table1) for different regions of EBV and normalized by the cellular DNA signal for the GAPDH or actin gene locus.

### Digital Droplet PCR

Digital droplet PCR was performed as previously described [32]. Briefly, DNA was isolated using the DNeasy Blood & Tissue Kit (Qiagen), and then DNA concentration was calculated using the NanoDrop2000 Spectrophotometer (ThermoFisher). DNA was digested and emulsified with droplet generator oil (Bio-Rad) using the QX-100 droplet generator and amplified using GeneAmp 9700 thermocycler (Applied Biosystems). The droplets were quantified using the QX200 droplet reader (Bio-Rad).

### Chromatin immunoprecipitation (ChIP)

For each ChIP assay, 1×10^6^ cells were crosslinked with 1% formaldehyde at room temperature for 10 min and the reaction was quenched with 0.125 M glycine for 5 min. Cells were lysed in 1 ml SDS lysis buffer (1% SDS, 10 mM EDTA, and 50 mM Tris-HCl, pH 8.0) containing protease inhibitor cocktails (Sigma-Aldrich), and kept on ice for 10 min. Lysates were sonicated with a Diagenode Bioruptor, cleared by centrifugation to remove insoluble materials, and diluted 10-fold into IP Buffer (0.01% SDS, 1.1% Triton X-100, 1.2mM EDTA, 16.7mM Tris pH 8.0, 167mM NaCl, 1 mM PMSF, and protease inhibitors cocktail). For each IP, 5 μg of anti-macroH2A (diagenode-C15210003), Anti-EBNA1 (Pocono Rabbit Farm custom preparation), anti-H2A.Z (Abcam-188314) or IgG (Cell Signaling, 2729), γH2AX (Abcam-ab243906) was added and rotated at 4°C overnight. Preblocked protein A sepharose (GE Healthcare, 17-0780-01/17-0618-01) was added to each IP reaction for additional 2 to 2 h incubation at 4°C. Beads were washed sequentially with low salt, high salt, LiCl, and TE buffer and then eluted with elution buffer (1% SDS, 0.1M NaHCO3). The eluates were then incubated at 65°C overnight to reverse cross-linking, followed by addition of Proteinase K at a final concentration of 100 μg/ml at 50°C for 2 hrs. ChIP DNA was purified by Quick PCR Purification Kit (Life Technologies) following the manufacturer’s instruction. ChIP DNA was assayed by qPCR using primers specific for indicated regions (**Table 1**). The relative enrichment was calculated as a percentage of input.

### Chromatin immunoprecipitation sequencing (ChIP-seq)

Chromatin immunoprecipitation with next-generation sequencing (ChIP-seq) was performed as previously described [12]. Briefly, 25×10^6^ cells per immunoprecipitation were collected and fixed with 1% formaldehyde for 15 min and then quenched with 0.25 M glycine for 5 min on ice. After 3 washes with 1× phosphate-buffered saline (PBS), pellets were resuspended in 10 mL each of a series of three lysis buffers before fragmentation in a Covaris ME220 sonicator (peak power 75, duty factor 25, cycles/burst 1,000, average power 18.8, time 720 s) to generate chromatin fragments roughly 200–500 bp in size as determined by DNA gel electrophoresis. Chromatin was centrifuged to clear debris and a 1:20 of this cleared chromatin was kept as standard input for comparison against immunoprecipitations. Chromatin was incubated by rotating at 4°C for 1 h with 25µg anti-anti-macroH2A (diagenode-C15210003). Chromatin–antibody complexes were precipitated using a 50 µL of Dynabeads Protein A (ThermoFisher, product No. 10001D) incubated by rotating at 4°C overnight. Beads were then washed five times with RIPA buffer and TE buffer. Crosslinks were reversed by 65°C incubation with proteinase K followed by RNase A treatment. DNA was purified by Quick PCR Purification Kit (Life Technologies) following the manufacturer’s instruction. Libraries for sequencing were made using the NEBNext Ultra II DNA Library Prep Kit (New England Biolabs, product No. E7103) and sequenced on the NextSeq 500 (Illumina).

### ChIP-seq analysis

Reads were mapped against the human genome and human gammaherpesvirus 4 (HHV4) NC_007605.1 genome assembly using bowtie2 [30]. We used MACS2 software packages to call reads enrichment in pull-down samples compared to input samples as peaks [31; 32]. Analysis of peak distribution under differentiated conditions was performed with the bedtools software package [33] for genome arithmetic, and for data visualization we used deepTools [34].

### Statistical analysis

Data showing the effects of different assays are presented as mean ± SD. p was calculated by Anova and Student’s t-test analysis. Significant differences were accepted with p ≤ 0.05, as indicated.

### Data Access

All sequencing data has been uploaded to NCBI GEO database under the accession number GSE353675 (RNAseq) and GSE353673 (ChIPseq).

## Results

### MacroH2A is enriched at the EBV oriLyt and terminal repeat regions

To determine if histone variant macroH2A associates with the EBV genome, we performed chromatin immunoprecipitation followed by sequencing (ChIP-seq) for macroH2A in two different EBV models, SNU-719 (EBVaGC-derived cell line) and Mutu I (EBV^+^ BL-derived cell lines (**Figure 1A-C**). We compared macroH2A ChIP-seq (blue) to Input controls (grey) and found enrichment at the repetitive DNA elements at OriLyt (l), BHF1, Orilyt(r), LF3 and TR (**Fig. 1B).** We also observed several narrow peaks at specific loci near BORF1, BALF2, and BARF1 (**Supplementary Figure 1**). Similar macroH2A ChIP-seq patterns were observed in SNU719 as in Mutu I. We confirmed by ChIP-qPCR that macroH2A was enriched at oriLyt, LF3, BHLF1 regions in SNU719 cells (**Fig. 1D**) and Mutu I (**Fig. 1E**). These results indicate that histone variant macroH2A is enriched in viral repetitive DNA elements including oriLyt, oriLyt associated transcripts BHLF1 and LF3, and the TR.

**Figure 1.**
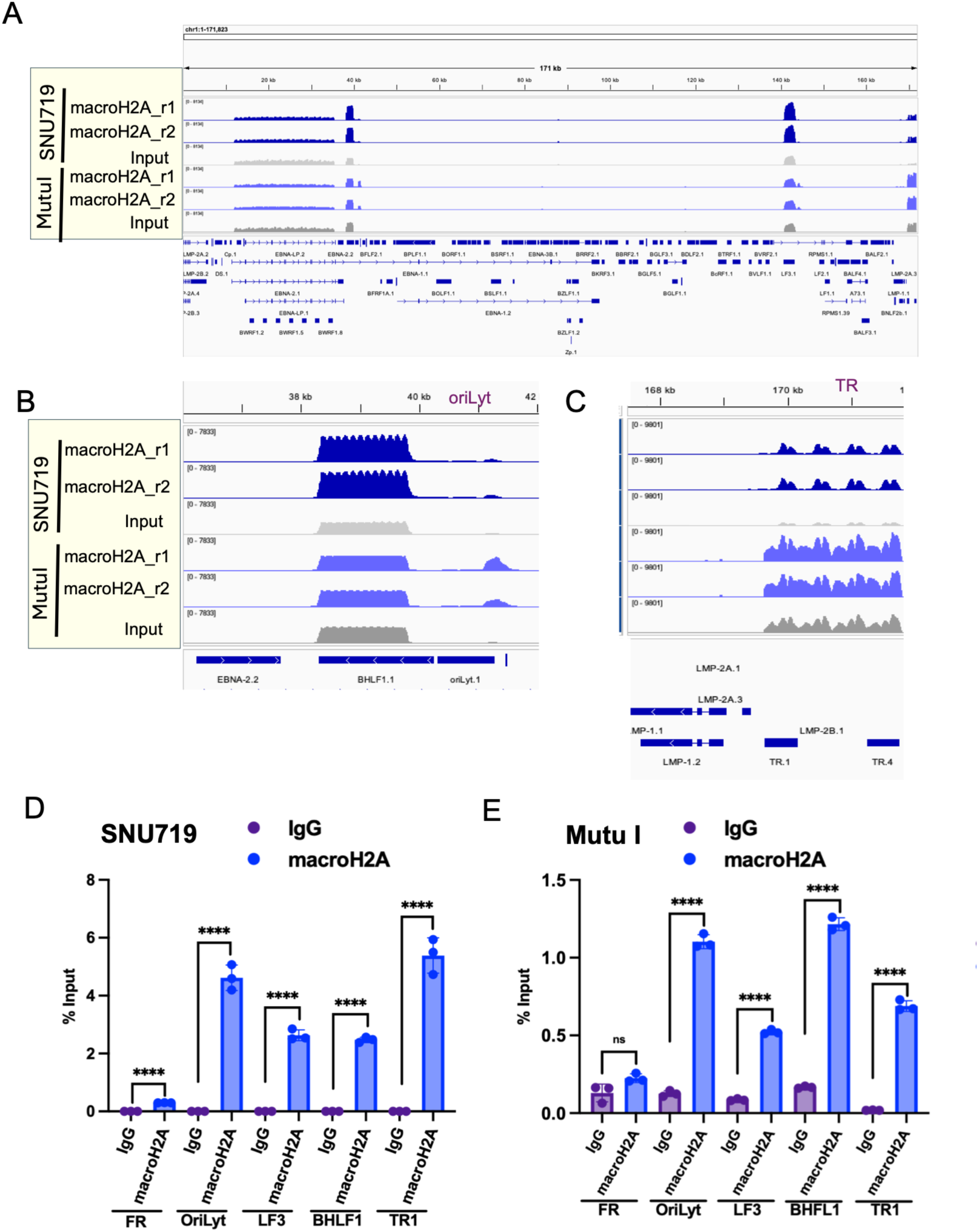
macroH2A is enriched at repetitive DNA and oriLyt in the EBV genome. (**A**) EBV full genome visualized using IGV showing ChIP-seq tracks for macroH2A (blue), and input control (grey) for SNU719 (top tracks) and Mutu I (lower tracks). (**B**) Magnified ChIP-seq tracks of panel A at BHLF1 and oriLyt. (**C**) Magnified ChIP-seq tracks for EBV TR. ChIP-qPCR of macroH2A or IgG control in SNU719 (**D**) and Mutu I (**E**) at FR, oriLyt, LF3, BHFL1 and TR1 regions of EBV. ****P < 0.0001, n = 3 independent experimental (ChIP) replicates, Student t-test.

### Macro-H2A knock-down alters transcriptional profile of SNU719

To gain insight into the function of macroH2A on EBV latency in SNU719, we performed siRNA depletion followed by RNA-seq. We first confirmed the siRNA pools to macroH2A.1 (*H2AFY*) efficiently depleted macroH2A by Western blot analysis (**Fig. 2A**). Western blot also showed that macroH2A depletion had no detectable effects on viral latency protein EBNA1 nor activation of immediate early lytic protein ZTA (**Fig. 2A**). Similar observations were made in other EBV-positive epithelial cancer cell lines, including gastric cancer cell line YCCEL1 and NPC cell line C666-1 (**Supplemental Figure 2**). Principal component analysis (PCA) revealed a clear separation of siControl and simacroH2A indicating that macroH2A depletion affects many significantly deregulated gene transcripts (**Fig. 2B**). Differential expression using DESeq2 (p-value of ≤0.05), identified 1,376 deregulated genes after the knockdown. Heat map with hierarchical clustering further demonstrated strong transcriptional response of simacroH2A and siControl (**Fig. 2C**). We identified 620 genes that were downregulated and 756 genes upregulated. Reactome Pathways analysis identified top upregulated pathways as Formation of Beta-Catenin TCF Transactivation Complex, Non-Integrin membrane ECM Interactions, Adipogenesis, and Chromatin Modifying Enzymes (**Fig. 2D**). The top-down regulated pathways including Formation of a Pool of Free 40S Subunits, Eukaryotic Translation Elongation, and Peptide Chain Elongation (**Fig. 2E**). These findings indicate that macroH2A depletion has little detectable effect on EBV proteins levels, but a significant impact on cellular transcriptome and differentiation state.

**Figure 2.**
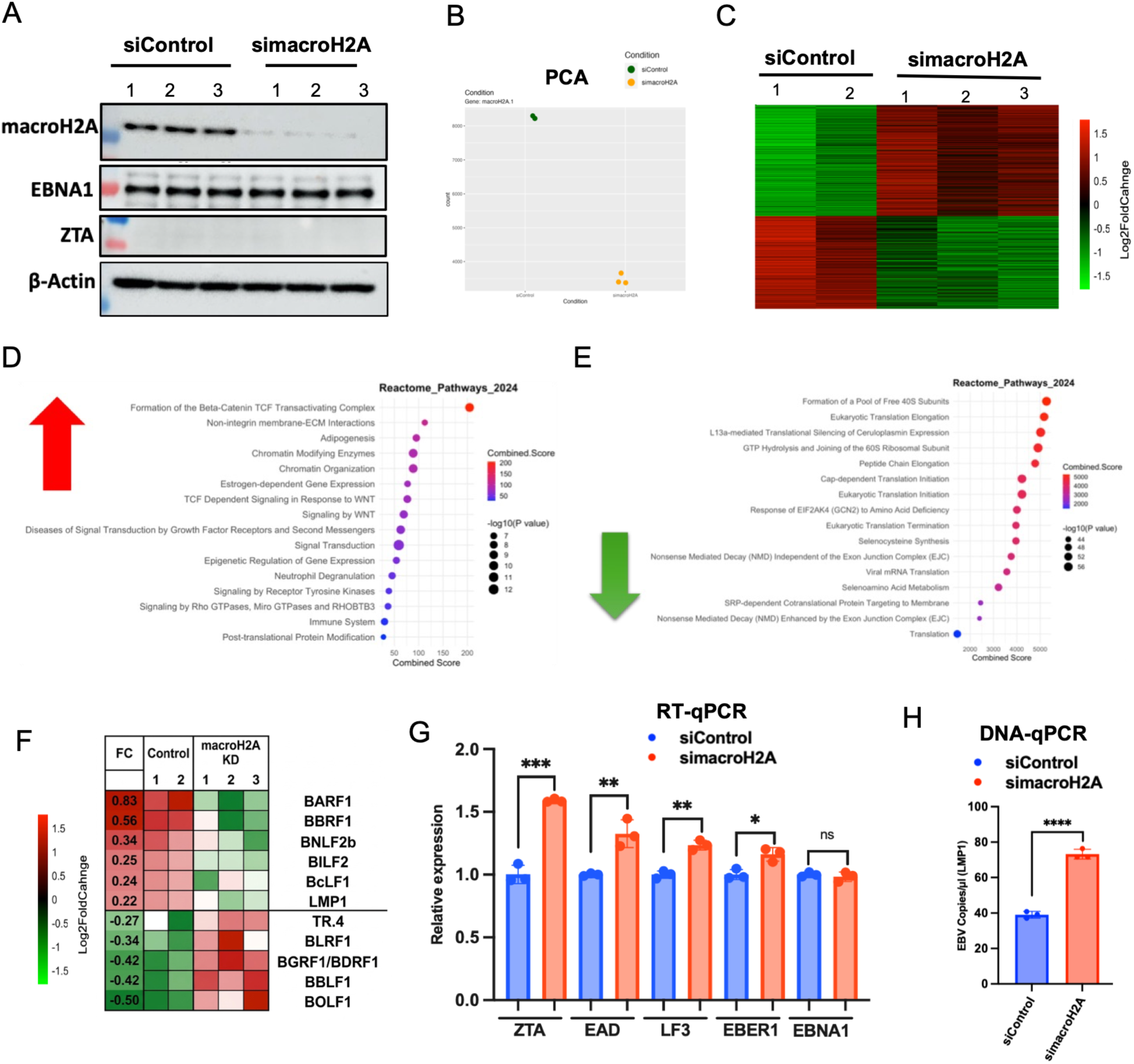
macroH2A knockdown affects host and viral transcriptomes. (**A**) SNU719 cells were transfected with siControl or simacroH2A and assayed by Western blot at 6 days post-transfection for expression of macroH2A, EBNA1, ZTA, or β-Actin. (**B**) Cells treated as described in panel A were subject to RNA-seq and analyzed by Principal Component Analysis (PCA). (**C**) Differential gene expression analysis was performed using DESeq2, and genes with statistically significant changes (P< 0.05) are shown by heatmap. (**D-E**) Pathway enrichment analysis of differentially expressed genes based on ShinyGO 0.8 analysis of the gene’s biological processes for top up regulated (**D**) or downregulated gene sets (**E**). (**F**) RNA-seq analysis of EBV transcripts from samples shown in panel A. (**G**) RT-qPCR for *ZTA, EAD, LF3, EBER1 and EBNA1* mRNA expression in SNU719 cells treated with siControl or simacroH2A. (**H**) EBV DNA copy number measured by Digital droplet PCR (ddPCR) using a probe for LMP1 gene relative to cellular RPP30 in SNU-719 transfected with siControl or simacroH2A. **P< 0.01, ***P< 0.001, ****P< 0.0001, Student t-test, n = 3 biological replicates.

### macroH2A knockdown has complex effects on EBV transcripts and DNA copy number

To determine if macroH2A regulates EBV transcript levels, we first analyzed RNA-seq data for changes in EBV genes (**Fig. 2F**). We found small effects on the viral transcriptome that included upregulation of some viral lytic cycle viral genes, including BLFR1, BGRF1/BDRF1, BOLF1, along with the terminal repeat transcript (TR.4). Downregulated viral genes including several latency associated transcripts, including BARF1, LMP2, BLF2b, as well as some transitional latent-lytic transcript such as BcLF1 and BILF2. We then assayed viral gene expression by RT-qPCR and found that macroH2A depletion led to a modest (< 2-fold) increase in EBV early lytic cycle transcripts, including *ZTA, EAD and LF3*, but also in some latency transcripts EBER1 (**Fig. 2G**). We next assayed whether macroH2A depletion had any effect on viral DNA copy number using DNA qPCR assay. We observed an ∼2-fold increase in EBV DNA copy number after macroH2A knockdown (**Fig. 2H**). Previous studies with knockdown of histone variant H2A.Z in the same SNU719 cells resulted in a greater increase (∼6 fold) in lytic transcripts and DNA copy number, as well as strong activation of ZTA and EA-D proteins detectable by Western blot [12]. Taken together, these findings suggest that while macroH2A depletion increased episome copy number, it does not induce full lytic transcription or replication.

### MacroH2A regulates EBV lytic gene transcription and DNA replication

Since was macroH2A was enriched at viral oriLyt regions by ChIP-seq and ChIP-qPCR (**Fig 1**) but had a minor effect on episome copy after knock-down in latently infected cells (**Fig. 2H**), we therefore tested whether macroH2A depletion had any effect on EBV lytic cycle replication after stimulation by an exogenous agent. We used 7.5 μM DCB for 3 days which is known to activate EBV lytic cycle efficiently by 48 hrs post treatment [35]. We validated that DCB induced the EBV lytic cycle immediate early protein ZTA ∼3.8 fold by 2 days post-treatment (**Fig. 3A-B**). We next tested the effects of siControl or simacroH2A on DCB induction of the EBV lytic cycle. We found that simacroH2A reduced the ZTA expression levels by ∼4 fold (**Fig. 3C-D**), although it did not prevent some activation above non-induced conditions. RT-qPCR analysis also showed that simacroH2A reduced transcript levels for *ZTA, EA-D and LF3*, relative to DCB treatment, but did not completely block transcript induction relative to DMSO (**Fig. 3E**). We next assayed DNA copy number by ddPCR and found that simacroH2A increased DNA copy number ∼2-fold relative to siControl+DCB, and ∼5 fold relative to DMSO (**Fig. 3F**). These findings suggest that macroH2A contributes a supportive activity for EBV lytic cycle transcription but has a restrictive activity for lytic DNA replication.

**Figure 3.**
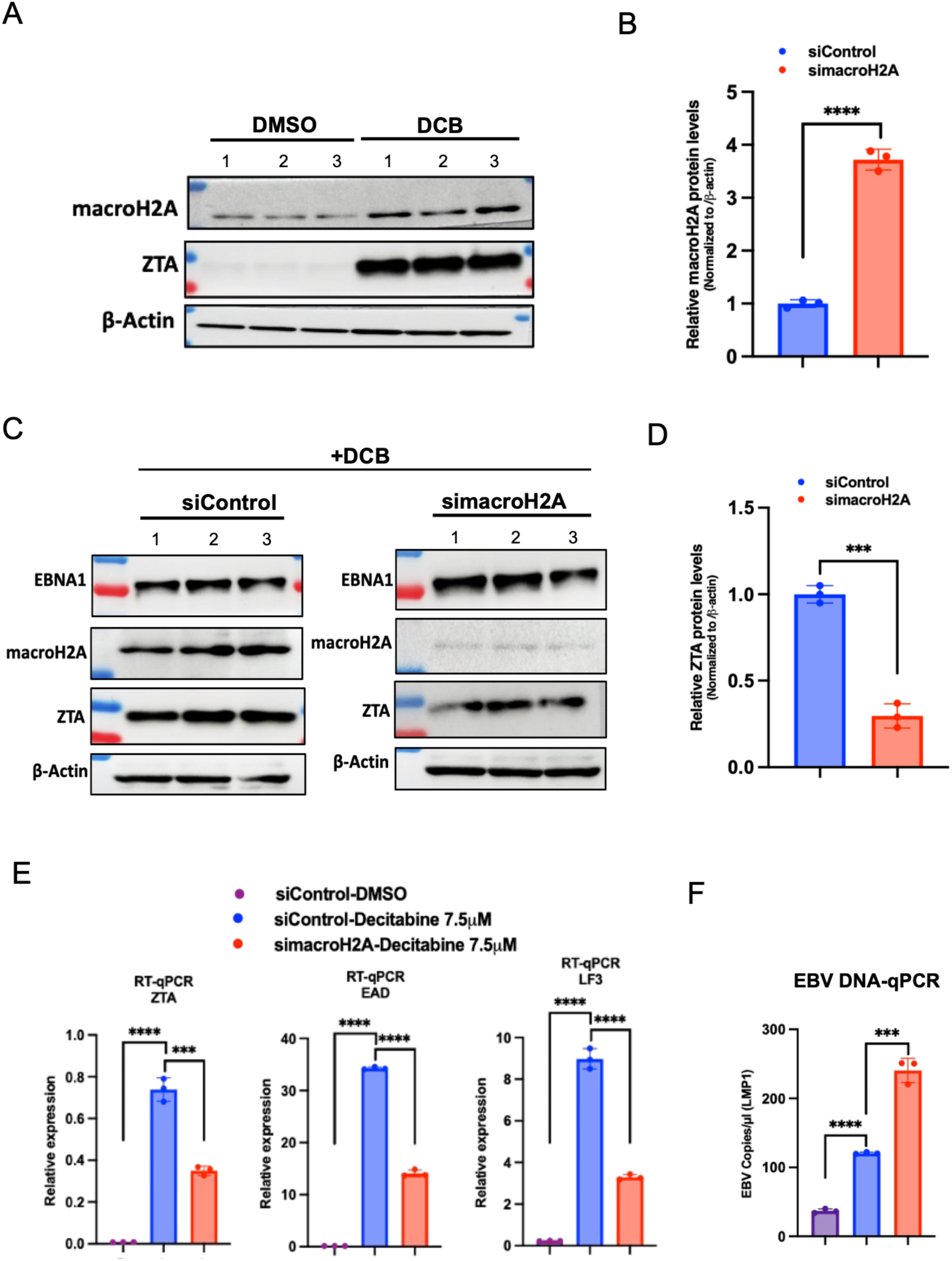
macroH2A histone variant participates in lytic cycle activation. (**A**) SNU719 cells. were treatment with 7.5µM of decitabine (DCB) and assayed by western blot for macroH2A, ZTA and β-actin. (**B**) Quantification of macroH2A protein relative to β-actin after decitabine treatment as shown in panel A. (**C**) SNU719 cells treated with 7.5µM of decitabine (DCB) were also treated with siControl or simacroH2A, and assayed by Western blot for EBNA1, macroH2A, ZTA, or β-Actin. (**D**) Quantification of ZTA protein levels shown in panel C. (**E**) RT-qPCR for *ZTA, EAD and LF3* mRNA expression in SNU719 cells treated with siControl+DMSO, siControl+DCB (7.5 μM) or simacroH2A+DCB (7.5 μM). (**F**) EBV DNA copy number measured by Digital droplet PCR (ddPCR) using a probe for LMP1 gene relative to cellular RPP30 as described in panel H. ****P< 0.0001, Student t-test, n = 3 biological replicates.

### Knock-down of macroH2A increases H2A.Z and EBNA1 occupancy at the EBV and cellular genome

Histone H2A variants may compete amongst each other for occupancy on chromatin. Our previous study found that histone variant H2A.Z colocalized with and facilitated binding of EBNA1 to EBV and cellular binding sites [12]. We therefore tested whether macroH2A depletion had any effect on H2A.Z or EBNA1 binding at known binding sites in the viral and host genomes (**Fig 4A**). ChIP-qPCR demonstrated macroH2A depletion increased H2A.Z binding FR (∼1.8 fold) and Qp (∼8 fold) (**Fig. 4B**). Similar increases in H2A.Z were observed at other known EBNA1 binding sites in the cellular genome, specifically SELK and CDC7 (**Fig. 4C**). We next assayed the effect of macroH2A depletion on EBNA1 binding at these same sites. Depletion of macroH2A led to a ∼5-fold increase in EBNA1 binding at FR and ∼2.2-fold increase at Qp (**Fig. 4D**). EBNA1 binding increased ∼8 fold at SELK and ∼4 fold at CDC7 (**Fig. 4E**). Together, these results indicate that the macroH2A depletion increases both H2A.Z and EBNA1 co-occupancies at viral and cellular binding sites.

**Figure 4.**
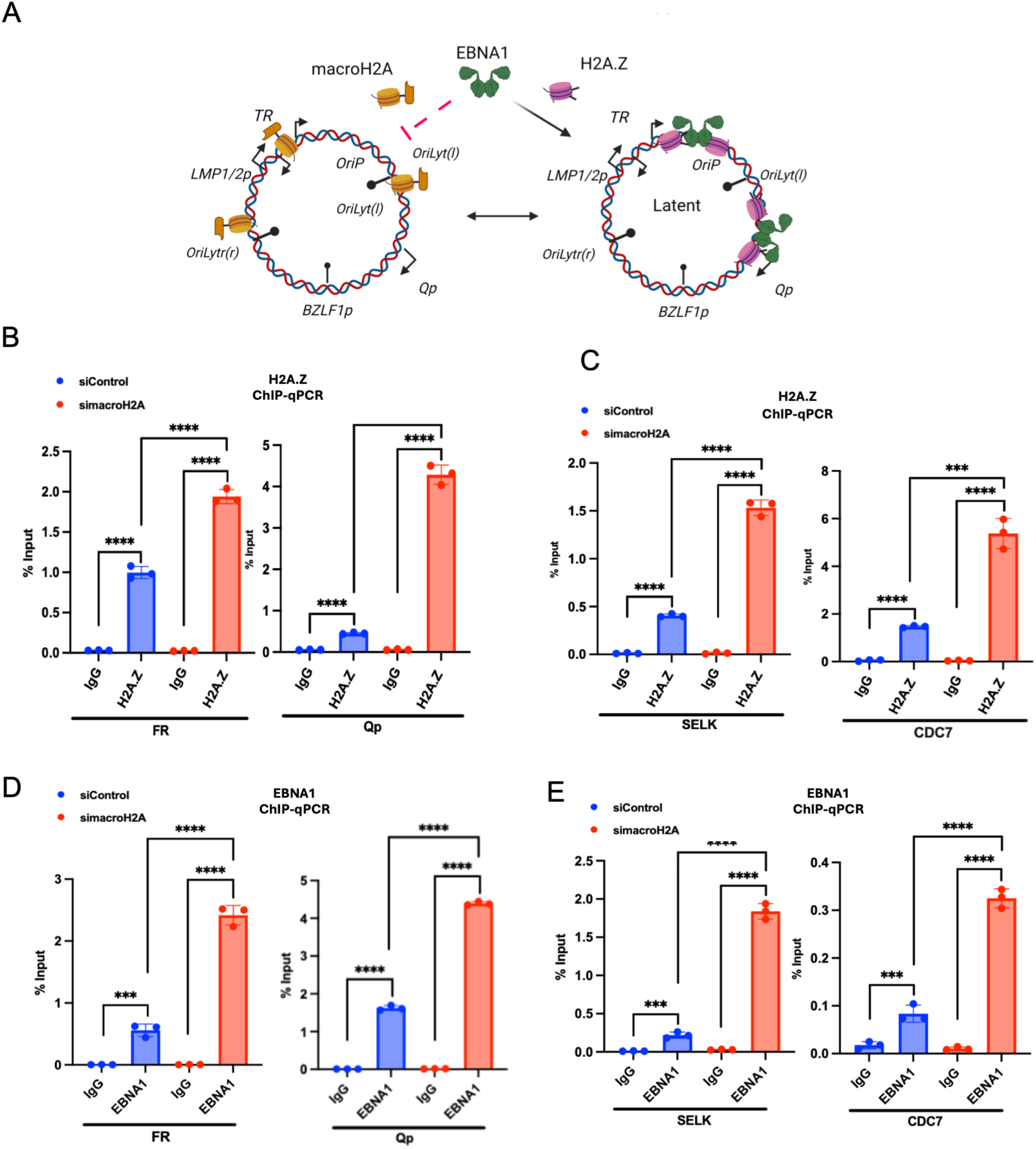
Knockdown of macroH2A increases EBNA1 and H2A.Z binding to the EBV genome and cellular genome. **(A)** Diagram of potential macroH2A and H2A.Z interplay on the EBV episome. (**B**) ChIP-qPCR for H2A.Z or IgG control at FR and Qp (**C**) ChIP-qPCR of H2A.Z for DNA binding at the cellular sites at SELK and CDC7 in SNU719 cells treated with siControl or simacroH2A. (**D**) ChIP-qPCR for EBNA1 or IgG control at FR and Qp. (**E**) ChIP-qPCR of EBNA1 for DNA binding at the cellular sites at SELK and CDC7 in SNU719 cells treated with siControl or simacroH2A. ****P< 0.0001, Student t-test, n = 3 biological replicates.

### macroH2A depletion decreases γH2A.X at oriP and induces changes in DNA damage response and DNA repair pathways

The histone H2A variant γH2AX localizes to sites of DNA damage and has been implicated in regulation of the EBNA1 orthologue LANA during KSHV latent cycle replication [36]. To investigate the potential role of γH2A.X at EBV oriP and how it may be affected by macroH2A depletion, we performed ChIP-qPCR in SNU719 treated with siControl or simacroH2A (**Fig. 5**). As expected, macroH2A depletion led to a loss of the macroH2A ChIP signal at LF3, oriLyt and TR1 (**Fig. 5A**). ChIP-qPCR for γH2AX identified FR and DS elements as highly enriched for γH2A.X (**Fig. 5B**). Interestingly, macroH2A depletion led to a significant loss of γH2AX binding at FR (∼4-fold reduction) and DS (∼10-fold reduction). These findings indicate the γH2A.X localizes to oriP, and that this localization depends on macroH2A.

**Figure 5.**
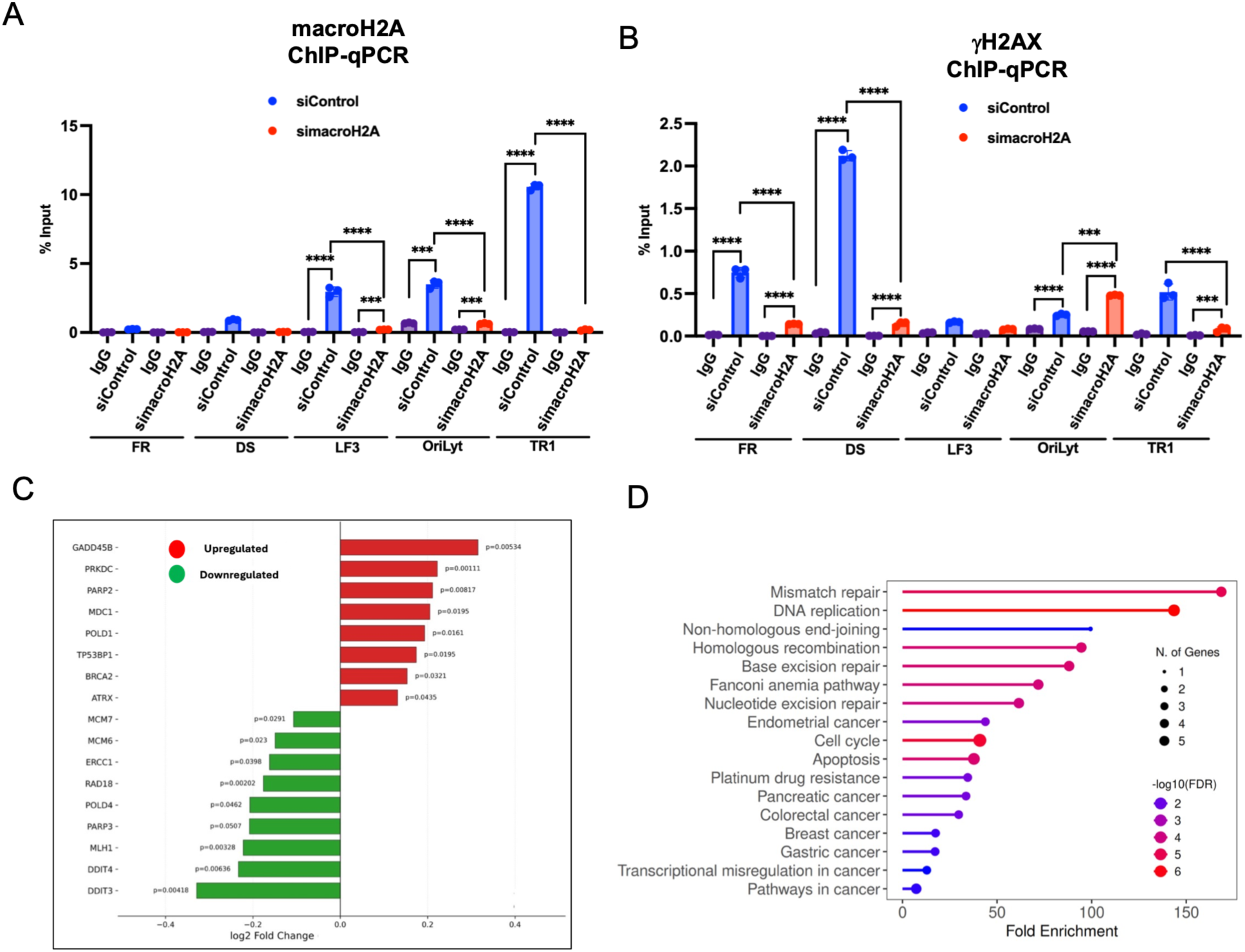
MacroH2A.1 knockdown induces changes in γH2A.X and DNA damage response and DNA repair pathways. **(A)** ChIP-qPCR for macroH2A or control IgG at the FR, DS, OriLyt and BHFL1 in SNU-719 cells treated with siControl and simacroH2A. (**B**) SNU719 cells were transfected with siControl or simacroH2A and assayed by ChIP-qPCR for γH2A.X or control IgG at the FR, DS, OriLyt and BHLF1. (**C**) Plot showing differentially expressed genes associated with γH2A.X from RNA-seq data after the knockdown macroH2A. (**D**) Pathway enrichment analysis of differentially expressed genes from panel C, based on ShinyGO 0.8 analysis of the gene’s biological processes. Differential gene expression analysis was performed using DESeq2, and all genes with statistically significant changes (P< 0.05). ****P< 0.0001, Student t-test, n = 3 biological replicates.

Since γH2AX is known to be a signal for DNA damage response, we further analyzed RNAseq data for DNA damage response genes affected by macroH2A depletion (**Fig. 5C and D**). We identified numerous DNA damage response genes that were both up regulated (GADD45B, PRKDC, PARP2, MDC1, POLD1) or down regulated (DDT3, DDT4, MLH1, PARP3, POLD4, RAD18). Pathways analysis showed a significant enrichment of Mismatch repair, DNA replication, and Homologous Recombination that were perturbed by macroH2A depletion in SNU719. These findings suggest that macroH2A plays a role in controlling the DNA damage associated with episomal EBV.

### Loss of macroH2A increases oriP-dependent plasmid DNA replication

To further investigate the role of macroH2A at oriP, we tested the effect of macroH2A knock-out on EBNA1-dependent oriP plasmid maintenance (**Fig. 6**). For these experiments, we utilized an EBV-negative EBVaGC cell line (AGS) with CRISPR/Cas9 targeted knockout of macroH2A (AGS sgmacroH2A) compared to control CRISPR/Cas9 AGS cells (AGS sgControl). Western blot demonstrated complete loss of macroH2A in both macroH2A sg1 and sg3, compared to sgControl AGS (**Fig. 6A and B**). These cell lines were then used for transfection with an oriP-containing plasmid expressing FLAG-EBNA1 (pOriP-EBNA1) or a control vector (pControl). Western blot confirmed the expression of EBNA1 in each of the transfected cell lines, although slightly lower levels were observed in the macroH2A knockout lines (**Fig. 6A and B**). Despite this lower level of EBNA1, we observed that macroH2A knockout cells had a ∼4.5-fold increase in oriP plasmid copy number compared with control (sgControl) (**Fig. 6C and 6D**). These findings demonstrate that macroH2A limits EBNA1-dependent oriP plasmid maintenance function.

**Figure 6.**
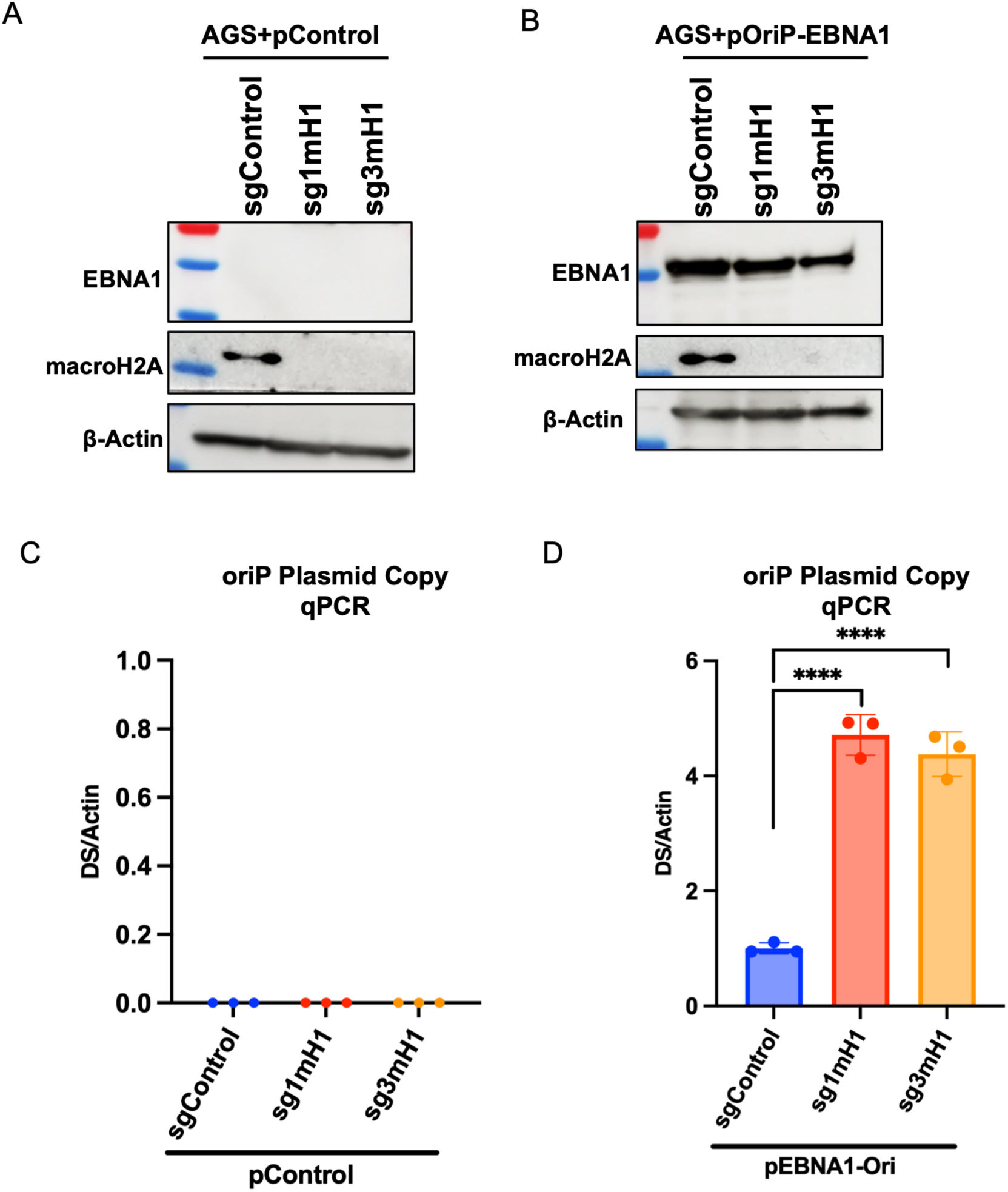
macroH2A knock-out increases oriP-dependent plasmid DNA replication in EBV negative cell line. **(A)** AGS cells with knockout of macroH2A.1 (one sgControl and two sg1macroH2A1 and sg3 macroH2A1) were transfected with control plasmid (pControl) and assayed by western blot for macroH2A, EBNA1 and β-actin. (**B**) AGS cells with knockout of macroH2A.1 (one sgControl and two sg1macroH2A1 and sg3 macroH2A1) were transfected with oriP plasmid expressing EBNA1 (EBNA1-OriP) and assayed by western blot for macroH2A, EBNA1 and β-actin. (**C**) OriP plasmid copy number determined by qPCR using primers for oriP elements DS region DNA for samples as described in panel A. (**D**) OriP plasmid copy number determined by qPCR using primers for oriP elements DS region DNA for samples as described in panel B. ****P< 0.0001, Student t-test.

## Discussion

EBV establishes latent infection as a circular, chromatinized episome that is regulated by various epigenetic mechanisms, including histone modifications and DNA methylation [37]. Histone variants are another epigenetic mechanism that can alter chromatin structure by modifying histone-protein or histone-DNA interactions, as well as the stability of nucleosomes that can affect gene expression and genome integrity [38]. To understand the role of histone variants in the regulation of the EBV genome, in this study we explored the role of the histone variant macroH2A, which is associated with heterochromatin architecture and genome integrity [39].

Global effects of macroH2A were examined by ChIP-seq and RNA-seq after siRNA depletion in SNU719 EBVaGC cell line. On the viral genome, we found that macroH2A is enriched at repetitive DNA regions of the EBV genome associated with lytic cycle regulation, such as OriLyt, BHLF1, LF3 and TR regions. Previous studies have found these regions subject to DNA and histone methylation linked to repression, such as H3K27me3, H3K9me2/me3, and H4K20me3 [35; 40; 41], consistent with their inactivity during latent infection. Using RNA-seq, we observed modest effects on viral transcripts, but significant effects on host genes and pathways. Depletion of macroH2A perturbs the expression of genes associated with several cellular processes including Beta-Catenin TCF Transactivation Complex, Non-Integrin membrane ECM Interactions, Adipogenesis, and Chromatin Modifying Enzymes. Many of these cellular processes have been reported previously and in other cell types to be regulated by macroH2A [24; 42; 43; 44; 45]. These cellular pathways have also been implicated in EBV infection and cancers. The β-catenin signaling pathway is downstream of PI3K activation which is frequently mutated in EBV epithelial cancers [46]. EBV is also implicated in EMT and invasiveness in epithelial cells [47; 48]. EBV latency transcription factors regulate chromatin modifiers [49; 50; 51; 52; 53]. Thus, macroH2A depletion modulates pathways known to be targeted by EBV.

The effects of macroH2A knockdown on EBV gene expression were small and limited to mostly lytic cycle transcripts such as BOLF1, BBLF1, BLRF1, and TR.4 (as determined by RNA-seq) and ZTA, EAD and LF3 (as determined by RT-qPCR). However, knockdown of macroH2A did not induce detectable lytic cycle proteins, including ZTA and EA-D, but did consistently increase EBV DNA copy number by ∼2 fold. This suggests that macroH2A depletion is insufficient to activate the full lytic cycle, but may promote a partial or abortive lytic response, in which the expression of selected viral genes is initiated but fails to progress toward a productive infection. EBV abortive lytic states have been described in several cellular contexts [54; 55]. Therefore, depletion of macroH2A may alter the epigenetic state of the episome that permits some lytic transcripts and episome amplification but fails to complete the full progression of the lytic cycle.

Consistent with a role of macroH2A in maintaining the balance between latent and lytic copy number control, we found that knockdown of macroH2A in cells treated with DCB leads to a decrease in lytic transcript, but an increase in viral copy number (Fig. 3). It is possible that macroH2A prevents the initiation of DNA replication during latency but supports lytic replication once it has been initiated. A potentially related observation was made by Lewis et al. (2023) demonstrating a functional role for macroH2A1 during herpes simplex virus (HSV) infection [56]. The authors showed that HSV-1 depletion of macroH2A1 or H3K27me3 resulted in reduced infectious viral titers and nuclear accumulation of viral capsids, despite having no significant effect on viral genome replication, viral RNA expression, or protein accumulation. They concluded that macroH2A1 dependent heterochromatin is essential for the translocation of HSV-1 capsids through the host chromatin to the inner nuclear membrane (INM), a step necessary for the release of viral particles [56]. Therefore, it will be of interest to further investigate the role of macroH2A during the EBV lytic cycle, as well as the occupancy and redistribution of macroH2A during from latency through the complete lytic replication and virion production cycle.

The interplay between macroH2A and other histone H2A variants was also observed. We previously showed that H2A.Z colocalizes with EBNA1 at oriP and other EBNA1 binding sites in the viral (e.g. Qp) and cellular genomes (e.g. SELK and CDC7) [12]. Our findings show that macroH2A depletion increases the occupancy of both EBNA1 and H2A.Z at viral and cellular genomic regions in SNU719 cells (Fig. 4). Furthermore, we observed an increase in the copy number of an oriP plasmid in AGS cells lacking macroH2A (via CRISPR/Cas9 knockout) that had been transfected with an EBNA1-oriP plasmid. These results suggest that macroH2A may compete with H2A.Z or otherwise prevent its assembly with EBNA1 and limited the formation of replication competent chromatin at oriP (Fig. 7). The effect of macroH2A on H2A.Z and EBNA1 binding could be indirect, but its effects can be observed on small episomal plasmids lacking other EBV transcripts, as demonstrated in the transient transfection experiments with EBNA1 and oriP plasmids (Fig. 5). Thus, it is more likely that macroH2A directly competes with H2A.Z for chromatin access and limits its binding at EBNA1 binding sites. The histone H2A variants have distinct effects on the structure and dynamics of nucleosomes [57] with some reports noting antagonism and opposing dynamics between them [58; 59], so it remains possible they act in a directly antagonistic manner and reciprocal replacement between the two occurs within EBV chromatin.

**Figure 7.**
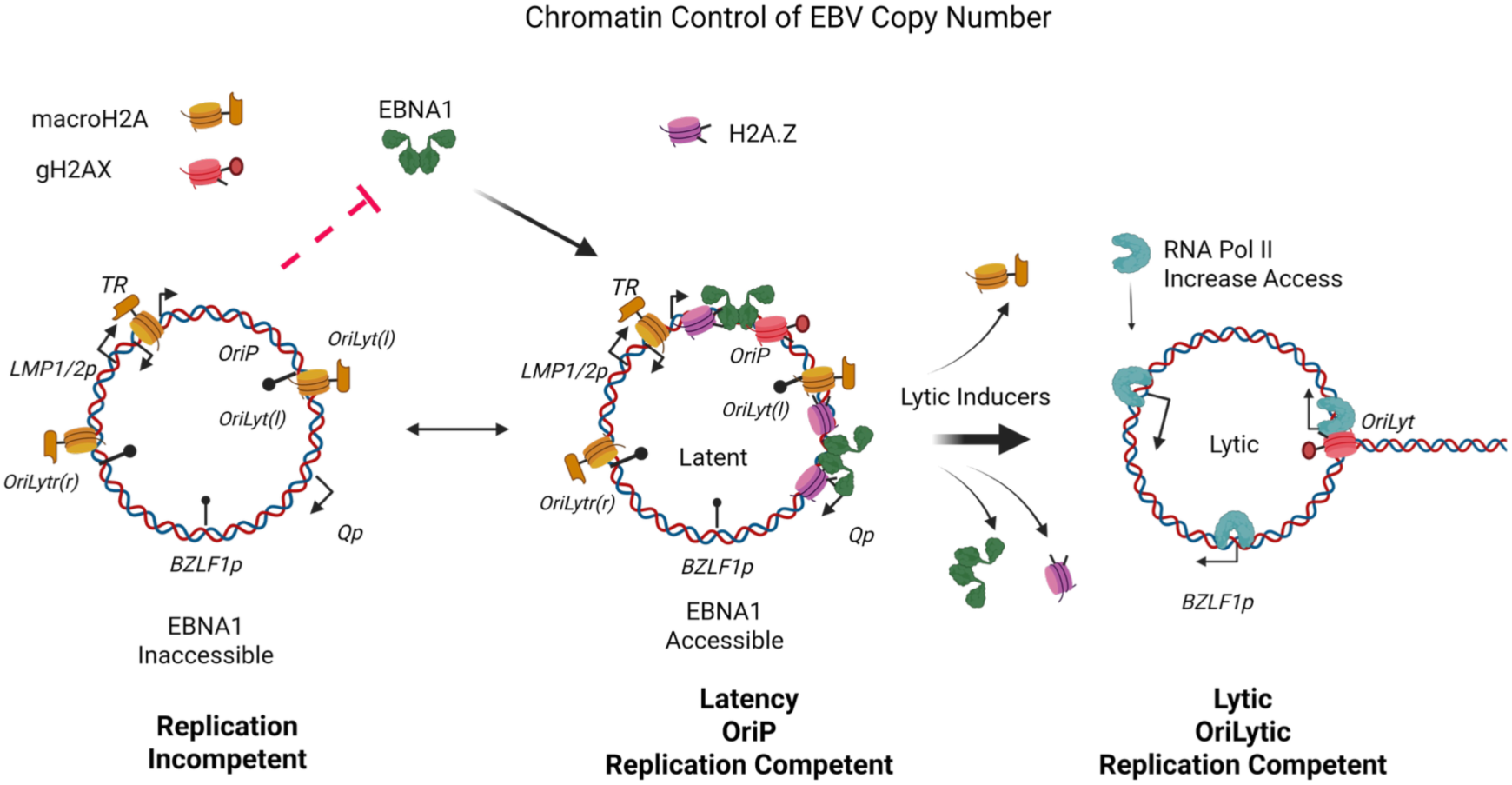
Model of histone H2A variant interplay in regulation of EBV episome maintenance. macroH2A antagonizes H2A.Z and EBNA1 binding, but promotes γH2AX formation at oriP.

Finally, we also found that another histone H2A variant γH2A.X was enriched at oriP and that macroH2A depletion reduced γH2A.X accumulation at oriP (Fig. 6). While γH2AX has been reported to bind to KSHV TR in association with LANA [36], it has not been previously shown to bind to EBV oriP during latent cycle replication. γH2AX is known to accumulate upon EBV and KSHV lytic cycle due to the replication stress and double strand DNA break signaling associated with rapidly replicating viral DNA genomes [49; 60; 61; 62]. However, some EBV latency proteins, including EBNA1, have been shown to induce γH2AX associated with replication stress [63; 64] and EBNA1 has been implicated in DNA cross-linking and strand breaks at oriP [65]. Thus, macroH2A may contribute to the chromatin dynamics associated with the oriP replication cycle, preventing over-replication by facilitating γH2AX recruitment and DNA damage repair at oriP. Consistent with this, macroH2A is essential for the stability and proper assembly of the machinery involved in the DNA damage or DNA stress response [13; 66], For example, macroH2A1.1 has been shown to facilitate the recruitment and retention of 53BP1 at sites of DNA double-strand breaks [67], while macroH2A1 contributes to BRCA1-dependent genome maintenance and the selection of DNA repair pathways [68]. Additionally, macroH2A1.1 regulates PARP-dependent signaling and promotes the efficient repair of oxidative DNA damage [18].

In conclusion, our findings identify macroH2A as an important regulator of EBV chromatin organization. Depletion of macroH2A altered the occupancy of factors associated with viral chromatin including an increase in the binding of EBNA1 and H2A.Z and led to greater accumulation of EBV DNA. Furthermore, it reduced the enrichment of γH2A.X in the viral regions analyzed and altered the expression of multiple genes involved in signaling, replication, and DNA damage repair. Taken together, these findings suggest that macroH2A plays a role in coordinating EBV chromatin organization with cellular mechanisms dedicated to genome maintenance. Notably, the effects of macroH2A depletion on the expression of EBV lytic genes were dependent on the experimental context: while depletion alone was insufficient to induce a complete lytic cycle, it did affect the lytic response to treatment with decitabine. Collectively, these results suggest that macroH2A serves various functions in EBV biology, influencing viral chromatin organization, the accumulation of episomal DNA, DNA damage-related pathways, and the cellular response to the induction of the lytic cycle. Therefore, macroH2A does not merely function as a repressor within the EBV genome; rather, it may contribute to maintaining the balance between EBV latency, episome maintenance, and lytic reactivation.

### 2.1 Tables

## 4 Author Contributions

Leonardo Josué Castro-Muñoz, Conceptualization, Investigation, Methodology, writing original draft. Leena Yoon, Formal analysis, Methodology, Visualization. Bhanu Chandra Karisetty, Formal analysis, Methodology, Visualization. Aradhana Bharti, Methodology, Formal analysis. Italo Tempera, Formal analysis, Visualization, Funding acquisition. Samantha S. Soldan, Formal analysis, Methodology, Visualization. Paul Lieberman, Conceptualization, Funding acquisition, Supervision, Writing, original draft.

## 8 Acknowledgments

We thank members of the Wistar Cancer Center Cores in Genomics and Bioinformatics for their excellent technical support.

## Supplementary Figures

**Supplementary Figure 1.**
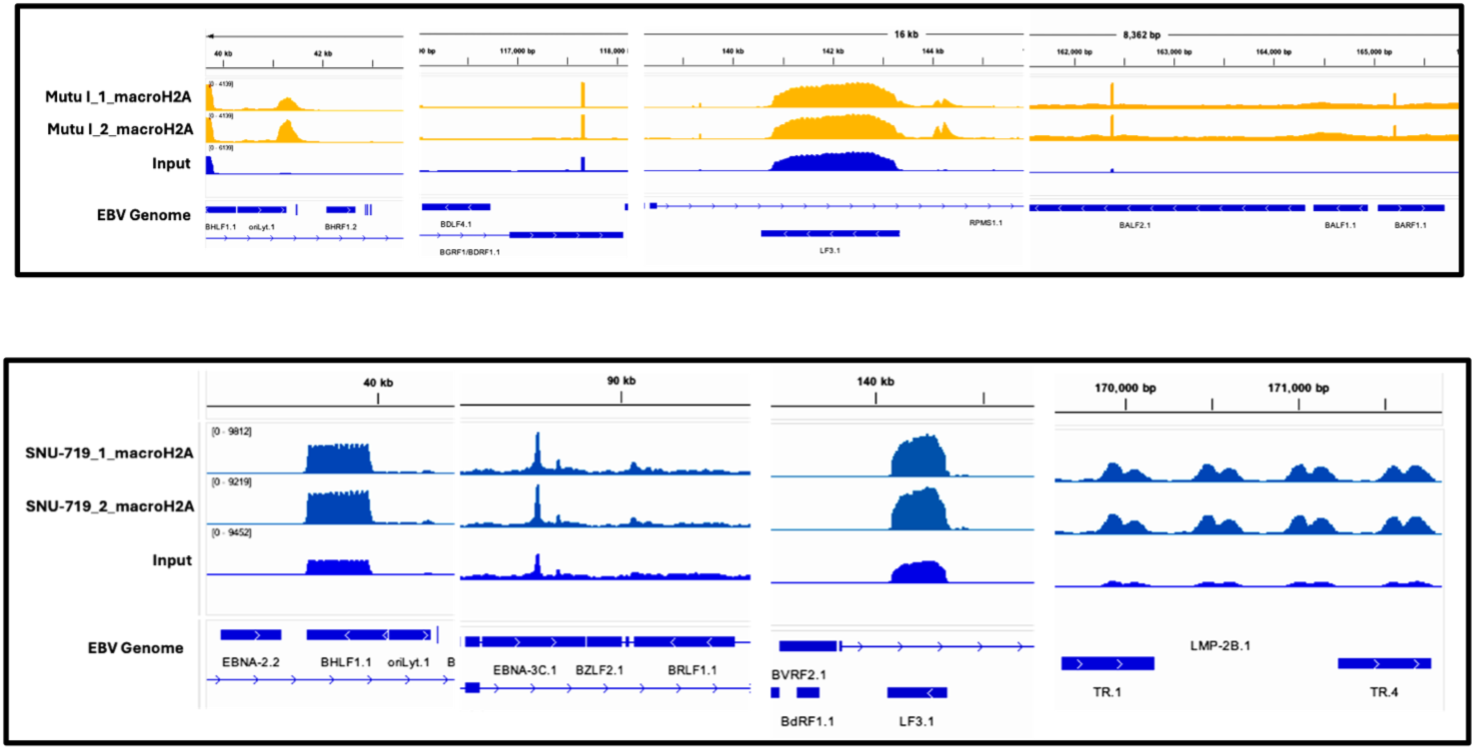
MacroH2A ChIP-seq narrow peaks. Narrow ChIP-Seq peaks at oriLyt, BGRF1, LF3, BALF2., BAR1.1 are shown in yellow for Mutu I (top) and at EBNA3C, LF3, and TR.1 for SNU71 (bottom).

**Supplementary Figure 2.**
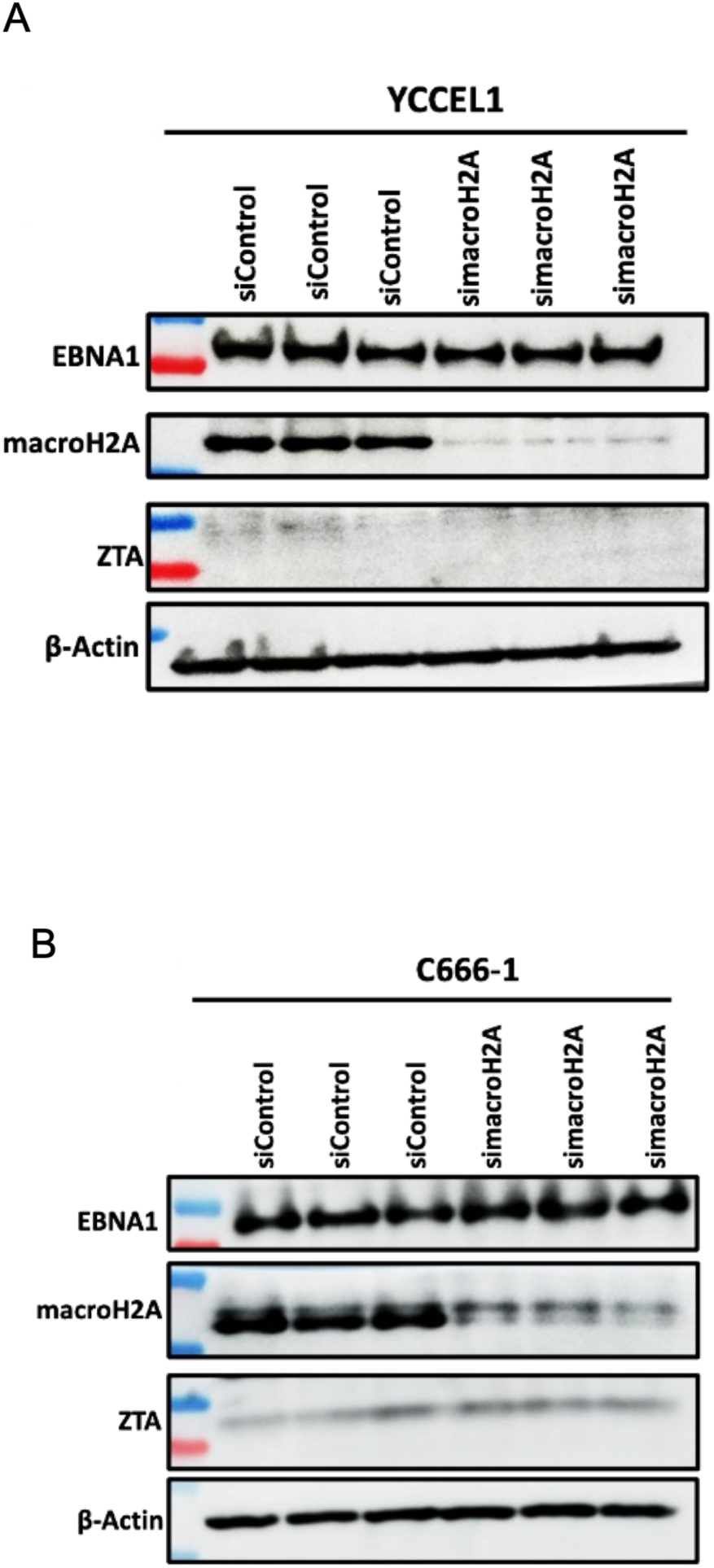
MacroH2A knock-down does not affect ZTA or EBNA1 protein expression in YCCEL1 or C666-1 cells. (A) YCCEL1 cells were treated with siControl or simacroH2A and assayed by Western blot for protein expression levels of EBNA1, macroH2A, ZTA or β-Actin. (B) C666-1 cells treated as described in panel A.

## 1 Data Availability Statement

All data is provided in the manuscript or supplementary materials. All raw sequencing data has been uploaded at NCBI GEO datasest under the accession number GSE353675 (RNAseq) and GSE353673 (ChIPseq).

